# Fenton-type chemistry by a copper enzyme: molecular mechanism of polysaccharide oxidative cleavage

**DOI:** 10.1101/097022

**Authors:** Bastien Bissaro, Åsmund K. Røhr, Morten Skaugen, Zarah Forsberg, Svein J. Horn, Gustav Vaaje-Kolstad, Vincent G.H. Eijsink

## Abstract

The discovery of Lytic Polysaccharide Monooxygenases (LPMOs) has been instrumental for the development of economically sustainable lignocellulose biorefineries. Despite the obvious importance of these exceptionally powerful redox enzymes, their mode of action remains enigmatic and their activity and stability under process conditions are hard to control. By using enzyme assays, mass spectrometry and experiments with labeled oxygen atoms, we show that H_2_O_2_, and not O_2_ as previously thought, is the co-substrate of LPMOs. By controlling H_2_O_2_ supply, stable reaction kinetics and high enzymatic rates are achieved, the LPMOs work under anaerobic conditions, and the need for adding stoichiometric amounts of reductants is alleviated. These results offer completely new perspectives regarding the mode of action of these unique mono-copper enzymes, the enzymatic conversion of biomass in Nature, and industrial biorefining.

**Abbreviations**
(AscA): ascorbic acid
(CDH): cellobiose dehydrogenase
(Chl): chlorophyllin
(GMC): glucose-methanol-choline oxidoreductase
(GH): glycoside hydrolase
(HAA): hydrogen atom abstraction
(LPMO): lytic polysaccharide monooxygenases
(pMMO): particulate methane monooxygenases
(O_2_^•−^): superoxide
(SOD): superoxide dismutase
(XTH): xanthine
(XOD): xanthine oxidase

---

The depolymerization of complex plant biomass, primarily composed of cellulose, various hemicelluloses and lignin, relies on a network of enzymatic and chemical reactions that is still full of mysteries. Until recently, the degradation of the recalcitrant polysaccharides in plant biomass was thought to be achieved by an arsenal of hydrolytic enzymes called glycoside hydrolases (GHs) (*1*). In some ecosystems, the enzymatic deconstruction process is supported by Fenton chemistry, i.e. transition metal-driven *in situ* generation of H_2_O_2_-derived hydroxyl radicals, one of the most powerful oxidizing species found on Earth (*2*), which can oxidize both polysaccharides and lignin in plant biomass (*3*). In 2010, a new class of enzymes was discovered, which carry out oxidative cleavage of polysaccharides (*4*). These enzymes, today known as lytic polysaccharide monooxygenases (LPMOs) (*5*), are single-copper redox enzymes (*6, 7*), that can hydroxylate the C1 or C4 positions of scissile glycosidic bonds (*4, 8–10*).

Despite their abundance in Nature (*5, 11*) and their obvious industrial importance, for example in the production of cellulosic ethanol (*12*), the mode of action of LPMOs remains enigmatic, although some catalytic mechanisms have been proposed (*7, 8, 10, 13, 14*). It is well-established that one LPMO reaction cycle requires the recruitment of two electrons (*4, 14–16*). The first electron is often thought to be acquired via reduction of the LPMO’s Cu(II) center to Cu(I) (*11*). When and how oxygen and the second electron are recruited remains an enigma. It appears impossible that an electron provider such as cellobiose dehydrogenase (CDH) (*11*) carries out direct reduction of the active site copper while the LPMO is bound to the substrate, whereas it is unlikely that the protein unbinds during catalysis to allow such a direct second reduction step. The existence of an internal electron channel that would allow electron delivery to a substrate-bound enzyme has therefore been postulated (*10, 17,18*).

Interestingly, a recent study has shown that unprecedented high levels of LPMO activity may be obtained when the enzyme is exposed to visible light in the presence of chlorophyllin (Chl) and ascorbic acid (AscA) (*19*). Although this study fell short of mechanistic explanations, the effect was attributed to the generation of high-energy electrons provided by photoexcited Chl, with AscA regenerating the Chl. From the increasing amount of publicly available data it appears clear that LPMO catalytic rates are indeed dependent on the nature of the redox partner (*11, 20*), which is intriguing, since it has been shown that the rate of LPMO reduction in solution is much higher (*11, 16*) than reported overall rates for LPMO action (*4, 14, 21*). These observations made us postulate that a chemical species, common to all known reaction systems but accumulating at different rates, plays an unsuspected key role in the LPMO mechanism. Looking for a potential culprit for LPMO activity, we studied the Chl/light, Chl/light-AscA and AscA systems for LPMO activation. A bacterial C1-specific cellulose-active LPMO10 from *Streptomyces coelicolor* (*Sc*LPMO10C) was used as primary model enzyme.

When using the Chl/light-AscA system, with relatively high light intensities, a strong increase in LPMO activity was indeed observed, notably accompanied by an almost immediate inactivation of the enzyme (**Fig. 1A**). Since light-exposed chlorophyll may produce superoxide (O_2_^•−^) (*22*), we investigated whether addition of superoxide dismutase (SOD) or superoxide-consuming chemicals to the Chl/light-AscA system would allow better control of the reaction, which turned out not to be the case (**Fig. S1**). On the other hand, we found that both the catalytic rate and apparent inactivation of the enzyme could be modulated by varying the amount of AscA (**Fig. 1A; Fig. S2**) or the light intensity (**Fig. S3**). Interestingly, in the absence of AscA, the Chl/light system yielded good LPMO activity and apparent inactivation of the enzyme was much reduced, as illustrated by a more linear progress curve for LPMO activity (**Fig. 1A**). Under these latter conditions, low concentrations of SOD were beneficial for LPMO activity, whereas high concentrations of SOD were detrimental due to rapid inactivation of the enzyme (**Fig. 1A; Fig. S4**). These results show that the levels of superoxide and/or the products of SOD, O_2_ and H_2_O_2_, affect LPMO activity.

**Figure 1C** shows that, in the absence of an LPMO, the Chl/light system produces H_2_O_2_ and that production is strongly increased by adding SOD, which enzymatically converts superoxide to H_2_O_2_, or AscA, which chemically reduces superoxide to H_2_O_2_ (**Fig. S5**). These H_2_O_2_ production levels in the absence of the LPMO (**Fig. 1C**) correlate well with the initial rates observed in the LPMO reactions (**Fig. 1A**). Moreover, rapid enzyme inactivation in the LPMO reactions (**Fig. 1A**) correlates with the H_2_O_2_ production potential (**Fig. 1C**) of the system used and is associated with accumulation of H_2_O_2_ in the reaction mixture (**Fig. 1B**). Notably, in the control reaction with only Chl/light, yielding relatively stable reaction kinetics (**Fig. 1A**), accumulation of H_2_O_2_ was not observed (**Fig. 1B**), whereas the Chl/light system does produce H_2_O_2_ (**Fig. 1C**). Addition of catalase reduced the detrimental effect of adding high amounts of SOD, reflected in slower inactivation of the LPMO (**Fig. 1A**) and reduced accumulation of H_2_O_2_ (**Fig. 1B**). All together, these results suggest that H_2_O_2_ is an unsuspected co-substrate for LPMOs and that too high levels of H_2_O_2_ are detrimental. The high initial LPMO rate observed when using the Chl/light+AscA system (**Fig. 1A**) is likely related to fast H_2_O_2_ production (up to 200 μM within the 12 first minutes of the reaction; **Fig. 1C**), which leads to rapid inactivation of the enzyme.

Control reactions with only AscA, well known for its ability to drive LPMO activity, yielded more modest H_2_O_2_ levels (< 40 μM within the first 60 min, **Fig. S6C**), which is likely related to AscA being less capable of engaging in the thermodynamically challenging reduction of O_2_, compared to Chl/light. Reactions similar to those in **Fig. 1** but only using AscA generally yielded less clear results (**Fig. S6A&B**), which is likely due to the many possible redox reactions involving AscA, superoxide and H_2_O_2_ (**Fig S5**). However, the same overall trend stood out: both higher LPMO activity and faster apparent enzyme inactivation were correlated with higher H_2_O_2_ levels.

**Fig. 1.**
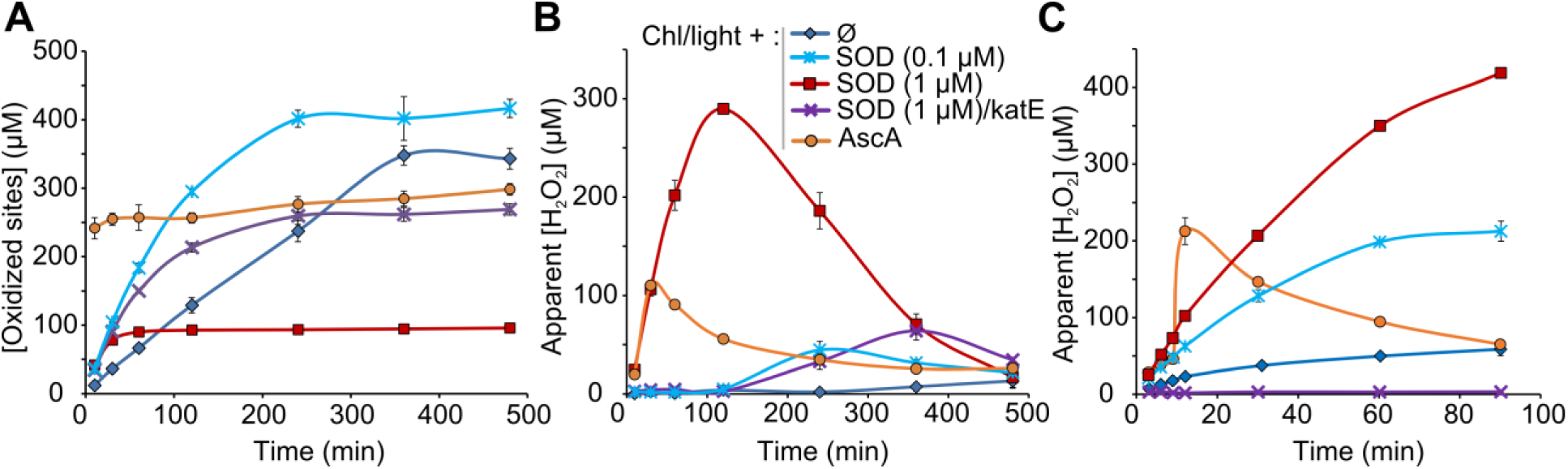
LPMO activity and apparent hydrogen peroxide production when using the Chl/light system for driving the reaction. Panels A and B show time-courses for the release of aldonic acid products (A) and H_2_O_2_ levels (B) upon incubating Avicel (10 g.L^-1^) with *Sc*LPMO10C (0.5 μM). Reactions were carried out in sodium phosphate buffer (50 mM, pH 7.0) at 40 °C, under magnetic stirring, with Chl (500 μM) exposed to visible light (I = 25% I_max_, approx. 42 W.cm^-2^). Note that, compared to the previous study by Cannella et al. (*19*), we used higher light intensities, which likely explains why in our hands, the Chl/light system also works in the absence of a reductant (blue diamonds). Reaction conditions varied in terms of the presence of SOD (0.1 or 1 μM), a catalase, katE (10 μg.mL^-1^), or AscA (1 mM). **Panel C** shows the apparent production of H_2_O_2_ in the absence of *Sc*LPMOl0C, with all other conditions being the same as for Panels A & B. Control reactions in the dark did not yield detectable levels of oxidized products (**Fig. S3A**), with the exception of reactions with AscA (**Fig. S6**). The legend code displayed in panel B applies also to panels A and C. Error bars show ± s.d. (n = 3).

Reactions with the Chl/light system (i.e. no AscA) seemingly lack a reductant needed to reduce the LPMO copper, which led us to speculate that O_2_^•−^ could be involved in LPMO reduction (**pathway (iv) Fig. S5**). Indeed, chemical (KO_2_) or enzymatic (xanthine/xanthine oxidase) O_2_^•−^ generating systems could drive LPMO activity, albeit at low levels (**Fig. S7**). Control experiments without any reductant but with exogenous H_2_O_2_ did not lead to cellulose oxidation (**Fig. S8**). This latter observation (**Fig. S8**) indicates that only the reduced LPMO can react with H_2_O_2_ and is crucial for the discussions below.

To determine the role of H_2_O_2_, we then analyzed initial LPMO rates in the presence of a reductant and varying concentrations of exogenous H_2_O_2_ (**Fig. S9**). A spectacular increase in initial LPMO rates was observed at the lower H_2_O_2_ concentrations, with up to 26-fold more soluble oxidized products being released from Avicel by *Sc*LPMOl0C after 2 minutes when incubated in the presence of 200 μM H_2_O_2_ (**Fig. S9C**). This increase in activity is in the same order of magnitude as the increases reported for the Chl/light+AscA system (**Fig. S3E** and (*19*)). At higher H_2_O_2_ concentrations, the LPMO reactions stopped very rapidly. Exogenous H_2_O_2_ affected the activity of a fungal LPMO9 from *Phanerochaete chrysosporium* K-3 (*Pc*LPMO9D) (**Fig. S9D-F**), another type of cellulose-active bacterial LPMO10, *Sc*LPMO10B (**Fig. S9G-I**), and a chitin-active LPMO10, CBP21 (**Fig. S9K-L**), in a similar manner, but significant differences were observed in terms of the degree of activity enhancement and the sensitivity to H_2_O_2_ (note that the rate enhancement for CBP21 is >100-fold; **Fig. S9L**) Control reactions in which the enzyme was replaced by Cu(II)SO_4_ did not show any oxidized products (**Fig. S10**).

The results described above suggest a catalytic mechanism in which an H_2_O_2_-derived oxygen atom, rather than an O_2_-derived oxygen atom, would be introduced into the polysaccharide chain. In the proposed mechanism (**Fig. 2**), a priming reduction of the LPMO-Cu(II) to LPMO-Cu(I) occurs first. H_2_O_2_ would then bind to the Cu(I) center and homolytic bond cleavage, similar to what happens during Fenton chemistry, would produce a hydroxyl radical. This likely leads to formation of a Cu(II)-hydroxide intermediate and a substrate radical by one of several possible pathways (**Fig. S11**). In each of these mechanisms, the reaction between a copper-hydroxyl intermediate and the substrate radical leads to hydroxylation of the substrate and to regeneration of the Cu(I) center, which can enter a new catalytic cycle.

**Fig. 2.**
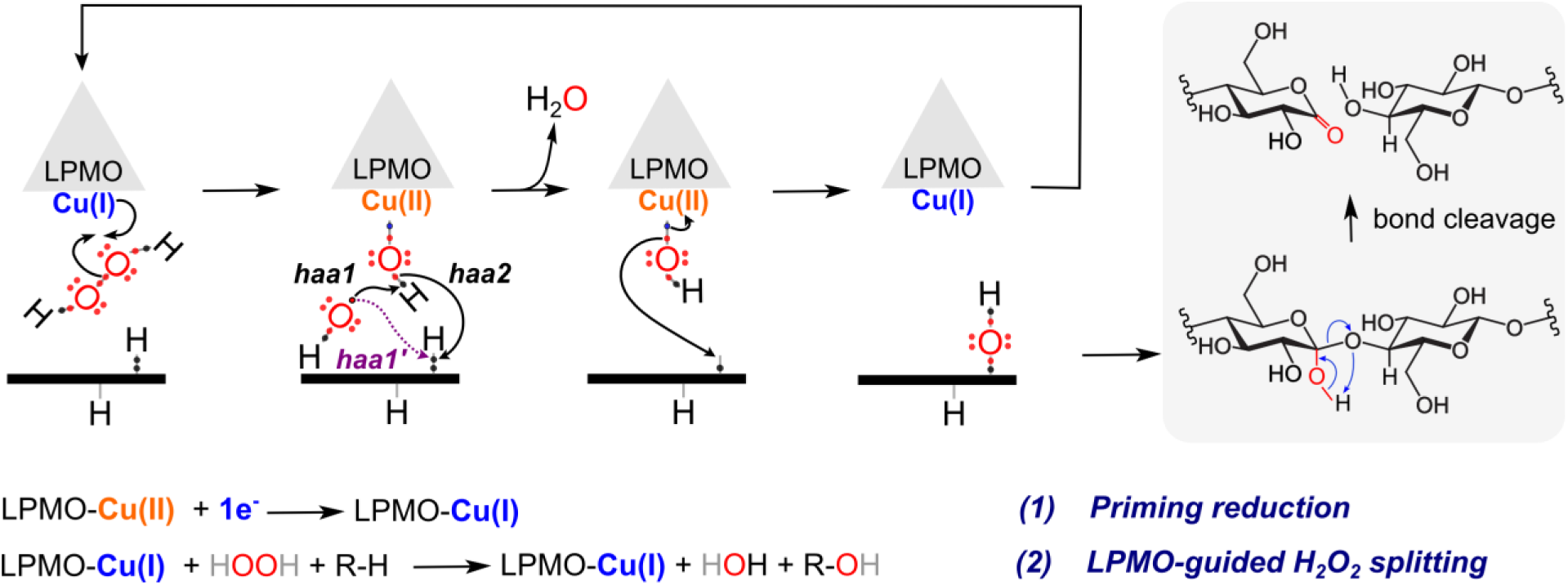
Proposed LPMO-guided H_2_O_2_ splitting mechanism for enzymatic oxidative cleavage of polysaccharides. In the proposed mechanism, LPMO-Cu(II) is first reduced to LPMO-Cu(I) (“priming reduction”), followed by H_2_O_2_ binding and homolytic bond cleavage. This cleavage leads to the Fenton-like generation of a hydroxyl radical, catalyzing HAA either from the Cu(II)-hydroxide (haa1) or from the substrate (haa1’). The former scenario would generate a copper-oxyl intermediate that can abstract a hydrogen atom from the substrate (haa2). In both scenarios a water molecule is eliminated and attack of the Cu(II)-hydroxide on the substrate radical leads to hydroxylation of the substrate and to regeneration of the Cu(I) center, which can enter a new catalytic cycle. The resulting hydroxylated polysaccharide undergoes molecular rearrangement leading to lactone formation and bond cleavage (*15*). The previously proposed reaction scheme for an O_2_-dependent reaction is: LPMO-Cu(II) + O_2_ + R-H + 2e^-^ + 2H^+^ → LPMO-Cu(II) + H_2_O + R-OH. See **Fig. S11** for further details.

To test this pathway and obtain final proof of H_2_O_2_ being the catalytically relevant co-substrate of LPMOs, additional experiments were carried out (**Fig. 3**). **Figures 3A&B** show that LPMO-dependent consumption of H_2_O_2_ (**Fig. 3A**) correlates with the release of oxidized products (**Fig. 3B**). Importantly, these experiments were done using catalytic (rather than putatively stoichiometric) amounts of reductant (10 μM; i.e. 100 times lower than commonly used concentrations; **Fig. S12**) to assess the concept of a “priming reduction” and to reduce the effect of AscA on H_2_O_2_ stability (**Fig. S13**). **Fig. 3B** shows that product levels are much higher than the total amount of AscA added. This is in agreement with the proposed mechanism in which a reduced LPMO can catalyze several reactions provided that the co-substrate, H_2_O_2_, is supplied. Analogous results were obtained when using a glucose oxidase from *Aspergillus niger* (*An*GOX), for controlled *in situ* generation of H_2_O_2_. **Fig. S14** shows that the glucose/AnGOX system boosts *Sc*LPMO10C activity in a dose-dependent manner, but only if the LPMO is reduced by a reductant added in small amounts (**Fig. S14**).

As a consequence of the above findings, LPMOs should be able to work under anaerobic conditions, which indeed was observed (**Fig. 3C**; **Fig. S15**). **Fig. 3C** shows that stable kinetics are obtained by adding H_2_O_2_ and reducing equivalents gradually to the reaction mixture and that the reaction rate is independent of the presence of O_2_. Finally, experiments with a labeled co-substrate, H_2_^18^O_2_, showed that indeed, the oxygen introduced into the polysaccharide chain comes from H_2_O_2_ and not from O_2_ (**Figs. 3D, S16-S18**). For example, **Fig. 3D** shows that when using H_2_^18^O_2_, the characteristic peaks for sodium adducts of the aldonic acid form of an oxidized cellohexaose *(m/z* 1029.7 &; 1051.7) shifted by +2 Da. Similar observations were made for the chitin-active LPMO10 CBP21 (**Fig. S16**) and a fungal cellulose-active LPMO9 (**Fig. S17**).Reactions with lower concentrations of H_2_^18^O_2_ showed that even in the presence of a 10-fold surplus of ^16^O_2_, the oxidized products carry ^18^O (**Fig. S18**). Finally, a competition experiment with a peroxidase and an LPMO showed that the peroxidase completely inhibited LPMO activity, despite the presence of O_2_ and reducing power (AscA or lactose/CDH; **Fig. S19**). Altogether, the experiments depicted in **Fig. 3** and **Figs S15-S19** unequivocally show that H_2_O_2_ is the catalytically relevant co-substrate for LPMO-catalyzed oxidation of a polysaccharide.

**Fig. 3.**
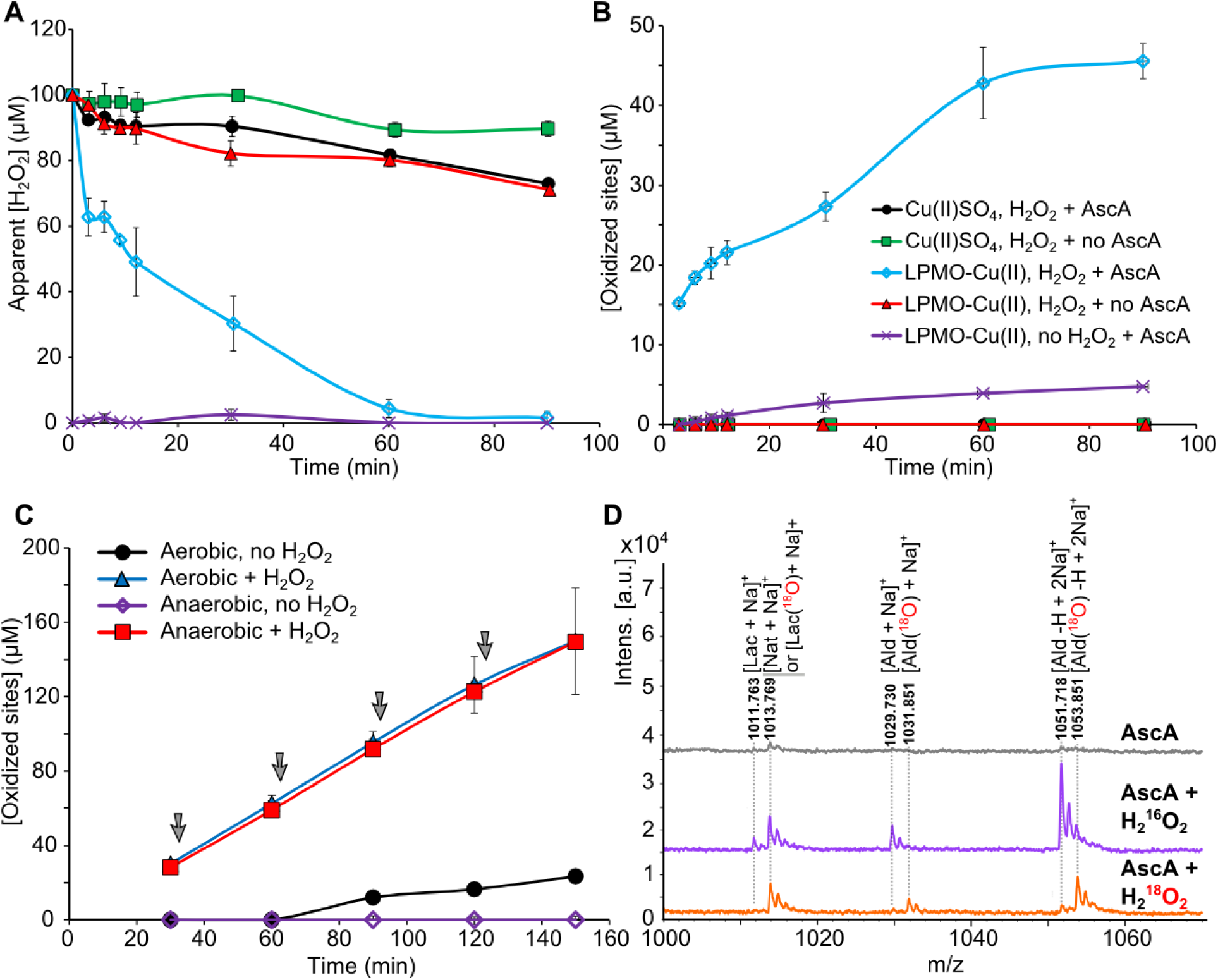
Probing the H_2_O_2_-dependent pathway of LPMOs. Panels A and B show H_2_O_2_ consumption (**A**) and soluble product formation (**B**) during incubation of *Sc*LPMO10C-Cu(II) and Avicel in the presence or absence of initial exogenous H_2_O_2_ (100 μM). In control reactions *sc*LPMO10C-Cu(II) was replaced by Cu(II)SO_4_ (0.5 μM). The reaction was initiated by addition of AscA (10 μM) where indicated. Note that this is a very low AscA concentration, meant to test the “priming reduction” hypothesis (see text for details). The legend code for panels A and B is indicated in the latter. Note that not all LPMO products are soluble, explaining why the levels of consumed H_2_O_2_ and detected products are not identical. (**C**) The graphs show time-courses for release of aldonic acid products by *sc*LPMO10C from Avicel under anaerobic or aerobic conditions at 30 °C. All the reactions were initiated by addition of AscA (10 μM) supplemented with H_2_O_2_ (50 μM) when indicated. AscA and H_2_O_2_ additions were repeated right after sampling, i.e. every 30 minutes (grey arrows). (**D**) MALDI-TOF MS spectra of products obtained after 4 min reaction in the presence of 100 μM H_2_^16^O_2_ or H_2_^18^O_2_, as indicated, and 1 mM AscA. The spectrum shows the hexose cluster, showing sodium adducts of the native (Nat) hexose, and the two forms of the oxidized hexose, the lactone (Lac) and the aldonic acid (Ald). All reactions (panel A to D) were carried out with *Sc*LPMO10C-Cu(II) (0.5 μM) and Avicel (10 g.L^-1^) in sodium phosphate buffer (50 mM, pH 7.0) at 40 °C under magnetic stirring (unless stated otherwise). The error bars show s.d. (n =3).

Several of the reaction progress curves discussed above show that LPMOs are readily inactivated and under some conditions, such as when using the Chl/light-AscA system (**Fig. 1A, Fig. S3D**), inactivation seems to occur within a few minutes. Enzyme inactivation was confirmed by a series of experiments where the LPMO was pre-incubated and then tested for remaining activity (**Fig. S20**). Enzyme inactivation was similar in the presence of EDTA, showing that inactivation is not due to free metal-catalyzed generation of hydroxyl radicals. Importantly, inactivation was partly avoided by the presence of substrate (**Fig. S20**). Using proteomics technologies, we found that the inactivated LPMO had undergone several oxidative modifications that were confined to the catalytic histidines and, to a lesser extent, neighboring residues (**Figs. 4, S21, S22**). Other residues prone to oxidative damage, such as surface exposed residues in the LPMO domain, the linker or the CBM were not modified (**Fig. S23**). This leads to the important conclusion that oxidative damage is not caused by ROS in solution, as has been suggested (*23*), but by ROS generated in the catalytic center, i.e. *in situ*, by enzyme-generated hydroxyl radicals with diffusion-limited timescale reactivity. The protective effect of the substrate (**Fig. S20, S24**), was reflected in reduced oxidative damage of the N-terminal catalytic histidine (**Fig. 4B**). The higher sensitivity of the N-terminal histidine may be related to the orientation of the reactive oxygen species during catalysis.

**Fig. 4.**
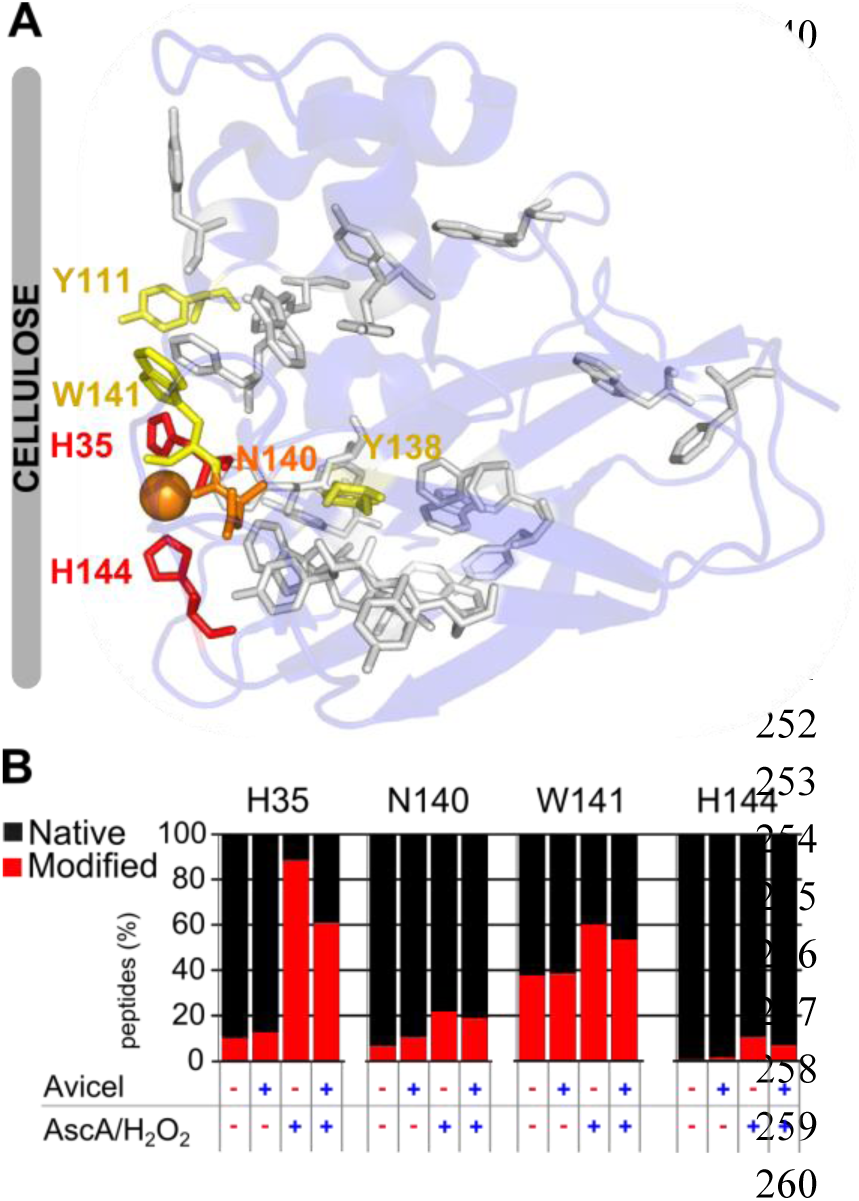
LPMO self-oxidation and the protective role of the substrate. (**A**) Mapping of modified residues on the structure of the catalytic domain of *Sc*LPMO10C (PDB 4OY7 (*24*)) reveals that oxidation occurs in and near the active site, predominantly on the catalytic histidines, H35 and H144. The color code highlights the degree of oxidation: high (red), middle (orange) and low (yellow). For aromatic residues shown as grey sticks no modification was detected (See **Fig. S21-S23**). (**B**) Impact of substrate on the ratio of modified/native peptides bearing H35, N140, W141 or H144 after a short incubation (i.e. a less drastic treatment compared to panel A). *Sc*LPMO10C (1 μM) was pre-treated by 20 min incubation in sodium phosphate buffer (50 mM, pH 7.0) at 40 °C under magnetic stirring, in the presence (10 g.L^-1^) or absence of Avicel and addition of either AscA (1mM)/H_2_O_2_ (100 μM) or simply water (control reaction). (See **Fig. S24** for corresponding activity tests).

The present findings unequivocally show that H_2_O_2_, and not O_2_, is the catalytically relevant co-substrate of LPMOs, implying that they should perhaps be called peroxygenases (or LPPO). Basically, LPMOs, after a priming reduction, carry out Fenton-type chemistry in a controlled and substrate-associated manner. From a biological point of view, such a scenario makes more sense than the seemingly somewhat random generation of dangerous hydroxyl radicals in the classical Fenton concept. Although the use of H_2_O_2_ by redox enzymes (e.g. peroxidases) is well-known, to the best of our knowledge, the biochemistry of the LPMO reaction as unraveled in this study, is unprecedented in Nature (*25, 26*).

LPMOs were initially classified as monooxygenases because experiments with ^18^O_2_ and using AscA as reductant, showed that one labeled oxygen atom was incorporated in the oxidized product (*4*). The proposed monooxygenase mechanism is well known (*26, 27*), seems “logical”, and has not been challenged until now. Strictly spoken, however, the seminal 2010 experiment with labeled oxygen did not show that O_2_ is the catalytically relevant co-substrate. In fact, other reactive oxygen species generated from O_2_ may have been the co-substrate, including H_2_O_2_, which indeed is produced under the conditions used (as shown in Fig. S6C). Importantly, the H_2_O_2_-pathway described here should not be confused with the “peroxide shunt” pathway that has been described for several monooxygenase reactions carried out by binuclear iron/copper enzymes, non-coupled binuclear copper enzymes and mononuclear iron enzymes containing additional co-factors such as porphyrin or biopterin. Such a “peroxide shunt” normally refers to a slow, rather artificial reaction that requires high concentrations of H_2_O_2_ (10-100 mM) and that is sometimes harnessed to avoid the use of reductants and O_2_ (*28, 29*). These shunt pathways involve the oxidized resting state of the enzyme and tend to lead to unstable reactions with a limited number of turnovers. The situation for LPMOs is very different and truly unique. LPMOs are mono-copper enzymes with no other co-factors, that, after an essential priming reduction, display stable reaction kinetics with multiple turnovers at low (sub mM) H_2_O_2_ concentrations.

Our findings explain several hitherto unexplained phenomena in LPMO biochemistry: (i) The consecutive delivery of two external electrons to the catalytic center is difficult to envisage, but with H_2_O_2_ being the co-substrate, recruitment of two electrons is not needed. (ii) The widely observed non-linearity of process kinetics is partly due to the self-inactivation of the LPMOs. (iii) The fact that most published catalytic rates for LPMOs are low and similar, and, most remarkably, independent of the LPMO or the substrate used (*4, 14, 21*), is likely due to the fact that the rate-limiting factor in most experiments was H_2_O_2_ formation. This point is well illustrated by a recent study demonstrating that the rate of a chitin-active LPMO fueled by the lactose/CDH system and the rate of H_2_O_2_ production by the latter system (in the absence of an LPMO) are similar (*16*). (iv) The increase in LPMO rate observed by Cannella et al. in their study on light-activation of LPMOs is due to production of hydrogen peroxide, not to the generation of some sort of “high energy electron” (*19*). (v) The observation that dehydrogenases can drive LPMO activity while strict oxidases cannot ((*30*) & Fig. S14) is due to the fact that, while both these enzyme types can produce H_2_O_2_, only the former can reduce the LPMO (*11*).

As to the level of H_2_O_2_ under reaction conditions, it is important to note that, notwithstanding the current findings, LPMOs are capable of activating molecular oxygen, albeit at low apparent rate (*8, 31*). It is well known that LPMOs generate H_2_O_2_ in the absence of substrate, which leads to the remarkable conclusion that LPMOs can generate their own co-substrate from O_2_. This property may have biological implications since H_2_O_2_ generated by unbound enzymes (several LPMOs display low substrate-binding) may be used by the substrate-bound population to degrade the substrate, explaining why H_2_O_2_ production by LPMOs is not observed in the presence of substrate (*16, 32*). It is conceivable that H_2_O_2_ interacts more strongly with substrate-bound, reduced LPMOs, compared to LPMOs in solution, which would explain why low concentrations of exogenous H_2_O_2_ are beneficial for activity, whereas higher concentrations lead to self-destructive reactions on unbound enzymes. Notably, the assumption that substrate-affinity has an impact on H_2_O_2_ management and self-destruction by the LPMOs sheds new light on the role of the CBMs that are appended to some LPMOs, including *Sc*LPMO10C.

The link between H_2_O_2_, Fenton-type systems and enzymatic biomass depolymerization has been a matter of debate, controversy and investigations for several decades. The present findings reveal a novel role for H_2_O_2_. Glucose-methanol-choline (GMC) oxidoreductases are known H_2_O_2_ producers and, like LPMOs, abundant in fungal secretomes (*11*). Some GMC oxidoreductases can reduce LPMOs (*11, 30*), but their ability to produce H_2_O_2_, perhaps in a controlled manner, could be another important biological function, as suggested by our experiment showing that a reduced LPMO can be fueled by the H_2_O_2_ generating glucose/glucose oxidase system. Along the same line, a recent study of the secretome of *Aspergillus nidulans* grown on starches revealed co-secretion of LPMOs, catalase and H_2_O_2_-producing oxidoreductases (AA3, AA7) in the absence of known H2O2-consuming partners such as peroxidases (*33*). It is noteworthy that the present findings may also be relevant for understanding host-pathogen interactions since for instance LPMO-producing necrotrophic bacteria are known to benefit from H_2_O_2_ generated by the plant defense system (*34*).

The present findings will have far-reaching implications for the design of biorefining processes, including the production of cellulosic ethanol. LPMOs are important components of current commercial cellulase cocktails (*12*) but proper aeration and delivery of electrons at industrial scale are considered major challenges as is the instability of LPMOs. We show here that LPMO performance and stability can be controlled by regulating the supply of H_2_O_2_, a liquid, cheap and easy-to-handle industrial bulk chemical. We further show that LPMOs can act in the presence of only catalytic amounts of reductant, which abolishes reductant-induced undesirable redox side reactions, and in the absence of oxygen, which eliminates the need for aeration. So far, the application of LPMOs has likely been hampered by suboptimal process conditions and it seems evident that further process improvements may be achieved now that the role of H_2_O_2_ has been uncovered. Notably, overdosing LPMOs can be a problem, since lack of LPMO binding sites on the substrate may lead to LPMO inactivation. It is conceivable that careful balancing of LPMOs and hydrolytic enzymes (e.g. cellulases) is needed, with the cellulases “peeling off” LPMO-disrupted polymer chains from the substrate surface, thus exposing new LPMO binding sites. As to LPMO stability, it is interesting to note that one of the residues most vulnerable to oxidation, the N-terminal catalytic histidine, is methylated in fungal LPMOs. Perhaps this methylation helps protecting the fungal LPMOs from oxidative self-destruction.

In the six years after their discovery (*4*), the role of H_2_O_2_ in LPMO catalysis has been overlooked, despite intense worldwide research on these enzymes. It is tempting to speculate that a similar situation may exist for other enzymes, in particular for copper monooxygenases that are thought to require two electrons and molecular oxygen. It has not escaped our notice that the still enigmatic particulate methane monooxygenase (pMMO), whose active site bears some resemblance to LPMO active sites, displays LPMO-like H_2_O_2_-related features: it has been reported that production of H_2_O_2_ by pMMO is lower in presence of substrate (*35*) and that H_2_O_2_ binds to and can oxidize the pMMO active site (*36*). It is conceivable that the present findings have implications beyond understanding and optimizing the enzymatic conversion of recalcitrant polysaccharides by LPMOs.

## Acknowledgments

We thank Bjørge Westereng at NMBU, Ås and Mats Sandgren at SLU, Uppsala, Sweden for providing a sample of a purified recombinant fungal AA9 (*Pc*LPMO9D). We thank Jennifer Loose at NMBU, Ås for providing the catalase katE. B.B. has received the support of the EU in the framework of the Marie-Curie FP7 COFUND People Programme, through the award of an AgreenSkills fellowship (under grant agreement n° 267196). The postdoctoral fellowship of B.B. was also supported by the French Institut National de la Recherche Agronomique (INRA) [CJS]. This work was also supported by the Research Council of Norway through grants 214613, 240967, 243950 and 249865, and by the Vista programme of The Norwegian Academy of Science and Letters through grant 6510.

## Supplementary Materials

Materials and Methods

Figures S1-S24

References *(1-60)*

